# HIV inhibits Warburg metabolism in human macrophages infected with Mycobacterium tuberculosis

**DOI:** 10.1101/2025.05.09.653039

**Authors:** Kevin Brown, Aaron Walsh, Anjali S Yennemadi, Dearbhla M. Murphy, Sarah A. Connolly, Mary P O’Sullivan, Sharee Basdeo, Seonadh O’Leary, Gina Leisching, Joseph Keane

**Author notes:** Corresponding author: Gina Leisching.

## Abstract

Tuberculosis (TB)-associated mortality remains disproportionately high among people living with HIV (PLWH), with macrophage dysfunction representing a key mechanism of impaired host defence against Mycobacterium tuberculosis (Mtb) infection. Using the U1 chronically HIV-infected macrophage cell line model coupled with primary human monocyte-derived macrophages (MDMs) exposed to HIV-1 gp120, we systematically characterized immunometabolic perturbations during Mtb infection. Nanostring RNA analysis revealed that Mtb monoinfection upregulated glycolytic genes while suppressing oxidative phosphorylation (OXPHOS) transcripts, consistent with a Warburg-type metabolic shift. Conversely, HIV infection downregulated glycolytic enzymes and enhanced mitochondrial respiratory chain components. Coinfection studies demonstrated HIV-mediated suppression of Mtb-induced glycolytic reprogramming. Extracellular flux analysis demonstrated that gp120 exposure increased basal oxygen consumption rate while impairing spare respiratory capacity in Mtb-infected MDMs, effectively blocking the Warburg metabolic transition. Notably, gp120 concentrations equivalent to those observed in antiretroviral therapy (ART)-treated PLWH significantly disrupted metabolic plasticity and high-dose gp120 attenuated Mtb-induced TNF-α secretion.

**Importance:** This study provides mechanistic insight into HIV-associated susceptibility to TB by demonstrating that HIV-1 infection fundamentally alters macrophage immunometabolic responses to Mtb. We establish that HIV-1, through gp120-mediated signaling, subverts the critical glycolytic induction required for effective antimicrobial responses against Mtb. The persistence of this metabolic dysregulation at clinically relevant gp120 concentrations, comparable to those observed in virologically suppressed PLWH, suggests ongoing immunological vulnerability despite ART. These findings identify HIV-induced metabolic reprogramming as a potential contributor to the persistently elevated TB risk in ART-treated individuals and highlight macrophage immunometabolism as a promising therapeutic target for host-directed therapies in HIV/TB coinfection. The dissociation between metabolic and cytokine responses suggests complex, multifactorial mechanisms underlying HIV-associated impairment of anti-mycobacterial immunity, warranting further investigation into the molecular pathways connecting cellular metabolism and immune effector functions.

## Introduction

Tuberculosis (TB) remains the leading cause of death in people living with HIV (PLWH), with HIV infection conferring a substantially increased risk of developing active TB disease. Macrophages play a critical role in early control of Mycobacterium tuberculosis (Mtb) infection, orchestrating inflammatory responses and executing antimicrobial functions such as phagocytosis, phagosome maturation, and cytokine secretion. However, HIV infection is known to impair these essential macrophage functions, contributing to poorer TB outcomes in PLWH(1). Despite this, the metabolic consequences of HIV/Mtb co-infection in human macrophages remain poorly defined, especially given the close association between metabolism and immunity.

HIV preferentially infects metabolically active CD4^+^ T cells(2, 3), capitalising on maximal available resources for rapid and extensive viral turnover(4). CD4^+^ T cells with high mitochondrial biomass are highly susceptible to infection, independent of their activation or differentiation status (5). Given this metabolic targeting, it might be expected that HIV-infected macrophages would also exhibit a distinctive metabolic phenotype. However, macrophages differ fundamentally from CD4+ T cells in their metabolism, tissue residency, and role in viral persistence. Studies investigating macrophage metabolism during HIV infection are limited(5) with the most reproducible finding being increased reliance on glutamine metabolism(6–8). This reliance is consistent with observations that HIV-infected macrophages show reduced mitochondrial oxidative phosphorylation (OXPHOS) and accumulate lipid droplets(7). The latter reflects an enhanced demand for fatty acids and cholesterol to support virion assembly and envelopment, a metabolic shift also seen in HIV-infected CD4+ T cells (9). However, direct comparisons between HIV-infected CD4+ T cells and macrophages reveal striking differences. Whereas HIV induces a glycolytic profile in CD4+ T cells, HIV-infected U937 macrophages exhibit reduced glucose uptake and no significant change in glycolytic intermediates (10). In primary macrophages, metabolic dysfunction has also been linked to viral proteins such as HIV Viral protein r (Vpr), which drives expression of glycolytic (11) and TCA cycle genes (8), further highlighting HIV’s ability to perturb macrophage metabolism in ways distinct from T cells. Vpr itself is essential for efficient viral replication in macrophages (12). These metabolic changes occur alongside well-documented HIV-mediated defects in macrophage antimicrobial responses. HIV co-infection sets the macrophage up for failure of early Mtb containment (13) through a variety of mechanisms, including reduced TLR4-stimulated TNFα secretion (14), failure of TNFα-mediated apoptosis in human AM and U1 macrophages (15), impaired phagocytosis (16, 17), and inhibition of phagosome maturation (18) and phagolysosomal fusion (19, 20).

Despite these established defects, no studies have directly examined how HIV alters macrophage immunometabolism during co-infection with *M. tuberculosis* (13). Given the importance of metabolic reprogramming for effective macrophage responses to Mtb (5), this represents a critical knowledge gap. Furthermore, antiretroviral therapy (ART) itself has been linked to mitochondrial toxicity (21–23), further complicating the interpretation of ex vivo studies in PLWH. Thus, characterising the transcriptional and functional metabolic signature of Mtb/HIV co-infected human macrophages and defining whether concomitant HIV infection corrupts macrophage glycolytic and mitochondrial metabolic reprogramming during Mtb infection might help to explain some of the persisting Mtb-specific immune defects seen in ART-treated HIV (24–30).

This study therefore explores human macrophage immunometabolism in Mtb/HIV co-infection in order to i) characterise the metabolic transcriptional signatures of chronically HIV-infected and Mtb/HIV co-infected human macrophages, ii) determine whether HIV co-infection impairs Mtb-induced glycolytic reprogramming and/or Warburg metabolism, iii) assess the effect of HIV gp120 treatment on mitochondrial function in Mtb-infected human monocyte-derived macrophages (MDM), and lastly iv) determine if HIV gp120 treatment will impair downstream macrophage anti-tuberculous human MDM effector function, including cytokine secretion.

## Methods

### Cell culture

U937 cells were obtained from the American Type Culture Collection (ATCC) and chronically HIV-1-infected U1 cells were purchased from the NIH AIDS Reagent Program. Cell lines were grown in RPMI completed with 10% FBS at 37°C and 5% CO2 and subcultured every 72hr. For experiments, U937 and U1 cells were matured from their monocytic to macrophage phenotype with 100 nM phorbol 12-myristate 13-acetate (PMA) (Sigma-Aldrich®, Cat# P1585-1MG) for 48 h. PBMCs were isolated from buffy coats of healthy donors (obtained with informed consent from the Irish Blood Transfusion Services) using Lymphoprep (StemCell Technologies) density-gradient centrifugation. Following isolation, cells were washed and resuspended at a density of 2.5 × 10^6^ PBMCs/mL in RPMI medium (Gibco) supplemented with 10% AB human serum (Sigma-Aldrich). The cell suspension was then plated onto non-treated tissue culture plates (Costar) and maintained in a humidified incubator at 37°C with 5% CO_2_ for 7 days. Non-adherent cells were removed by washing every 2–3 days. MDM purity was assessed by flow cytometry and consistently exceeded 90%.

### M. tuberculosis infection

Virulent *Mycobacterium tuberculosis* (H37Rv) acquired from the ATCC was grown in Middlebrook 7H9 broth culture medium (Sparks Ltd), supplemented with 10% albumin-dextrose-catalase (ADC) (Sparks Ltd, Cat# 211887) in an upright Nunc® T25 culture flask and maintained at 37°C and 5% CO2 in the CL 3 facility incubator and grown until log-phase. Gamma-Irradiated Mycobacterium tuberculosis (iH37Rv) was acquired from Bei Resources, NIAID, NIH. Macrophages were infected with a low multiplicity of infection (MOI) represented as ~30% infectivity and 1-5 bacilli/cell as previously described(31).

### Macrophage stimulations

#### γ-irradiated iH37Rv Stimulation

All bioenergetic flux Mtb stimulations were performed using γ-irradiated iH37Rv. After determination of MOI, MDM were stimulated with the desired volume of iH37Rv either 30-60 minutes before bioenergetic flux analysis, or during the bioenergetic flux run by direct injection from the cartridge port. Both of these methodologies are described in detail in the “Bioenergetic Flux Analysis” section below.

#### HIV gp120 Stimulation of Human Macrophages

Recombinant HIV-1 IIIB Envelope Glycoprotein 120 (Baculovirus) (ImmunoDx®, Cat# 1001) was acquired from the Centre for AIDS Reagents (CFAR), National Institute for Biological Standards and Control (NIBSC) (Repository Reference EVA607). Previously seeded human MDM in plastic tissue culture plates/Agilent® Seahorse® 24-well XF Cell Culture Microplates were treated for 24 hr with HIV gp120 at either low (100 ng/mL) or high (5,000 ng/mL) dose and rested for 24hr at 37°C and 5% CO_2_. Low dose HIV gp120 was used to represent “ART-treated HIV infection”, as this concentration of HIV gp120 has been reported in BAL fluid of ART-treated PLWH (32). High dose HIV gp120 was selected to test whether any findings were dose-dependent, and as a comparative “untreated HIV” model (340). MDM cell culture medium was replaced, and freshly thawed HIV gp120 was diluted in complete RPMI cell culture medium to desired concentration (100ng/mL or 5,000ng/mL) for the experimental treatment groups. After 24 hr cell culture medium was removed, cells were gently washed with warm sterile PBS and fresh cell culture medium (without HIV gp120) was replaced.

#### Gene Expression and Transcriptional Analysis

Total RNA was harvested from human macrophages 24 hr following H37Rv Mtb infection and isolated and purified by silica-membrane column centrifugation, using the RNeasy® Plus Mini Kit (QIAGEN®). RNA was assessed using the NanoString® nCounter® Metabolic Pathways panel. Metabolic mRNA signatures for four comparable conditions were characterised: “Control” (uninfected U937 macrophages), “TB” (Mtb-infected U937 macrophages), “HIV” (Mtb-uninfected U1 macrophages) and “TB/HIV” (Mtb-infected U1 macrophages). Data were analysed using the NanoString® nSolver™ 4.0 software and Advanced Analysis module to generate raw mRNA counts for each condition, which were then normalised and expressed as log2FC values using the in-built housekeeping, normalisation, and QC processes. To control for multiple testing, p-value adjustment was performed using the BH method for calculation of FDR. All p-values presented are adjusted p-values (q-value), unless stated otherwise. Data visualisations were generated by the NanoString® nSolver™ 4.0 Advanced Analysis module and R programme (https://www.r-project.org).

#### Bioenergetic Flux Analysis

For the acute injection protocol, MDM culture medium in Agilent® Seahorse® 24-well XF Cell Culture Microplate was replaced with 500µL of supplemented warmed Seahorse® XF RPMI Medium, and cells were rested in the non-CO2 cell culture incubator at 37°C for 30-60 minutes. During this resting period, injection ports of the calibrated Agilent® Seahorse® XFe24/XF24 Sensor Cartridge were filled with pre-determined treatments. For the Mtb treatment wells, the desired volume of iH37Rv (as determined by MOI) was placed in Port A and topped up to a final volume of 56µL with completed Seahorse® XF RPMI Medium. For non Mtb treatment wells, and for the four “blank” wells, 56µL of completed Seahorse® XF RPMI Medium was placed in Port A. Ports B, C and D were left empty for all 24 wells and were not included in the injection protocol during the run. Mito Stress tests were performed using the Seahorse® XF Cell Mito Stress Test Kit (Agilent Technologies®), according to the manufacturer’s instructions. Dilutions for each of oligomycin, FCCP and Rotenone/Antimycin A was then made for injection concentrations of 10µM, 10µM and 5µM, respectively. After each run, MDMs in the microplate were fixed for normalisation using Crystal Violet Staining Solution (0.5%).

#### ELISA

Supernatants were assessed by ELISA for the detection of TNFα (Invitrogen, 88-734688), IL1β (Biolegend, 437004) and IL-10 (Biolegend, 430604) according to the manufacturer’s instructions.

### Statistics

Statistical analyses were performed using GraphPad Prism 9 software. Statistically significant differences were determined using Student’s paired t-tests with two-tailed P-values. Differences between three or more groups were determined by one-way ANOVA with Tukey’s multiple comparisons tests. P-values of <0.05 were considered statistically significant.

## Results

### Mtb, HIV and Mtb/HIV Co-infection Induced Markedly Distinct Transcriptional Profiles in Human Macrophages

Transcriptional signatures across each of the conditions were found to be significantly different from each other. As can be appreciated from the PCA plot in Figure 1A, there was clear separation between each of the treatment groups, indicating clear and consistent changes in gene expression. Those genes found to be most markedly differentially expressed between each of the conditions are shown in the heatmap in Figure 1B, where clear clustering of the treatment conditions was observed, reinforcing the notion that both Mtb and HIV infection induced significant gene expression changes. The volcano plots in Figures 1C, 1D and 1E demonstrate the magnitude of gene expression changes in the “Mtb”, “HIV” and “Mtb/HIV” infected groups, respectively. Those genes found to be most markedly differentially expressed clustered into several important gene sets, and are shown in Table 1. Mtb generally induced genes within the “Cytokine and Chemokine Signalling”, “NFκ B” and “Tryptophan/Kynurenine Metabolism” gene sets, including the most upregulated genes IL-6, SOD2 and IDO1 vs. control (Fig.2). HIV infection, on the other hand, was associated with reduced expression of genes within the gene sets “Cytokine and Chemokine Signalling”, “Tryptophan/Kynurenine Metabolism” and “Glycolysis”. Mtb/HIV co-infected macrophages exhibited a mixed transcriptional profile with impaired “Cytokine and Chemokine Signalling” induction, and differential gene expression profiles within “Glutamine Metabolism” and “Glycolysis” gene sets compared to Mtb-infected macrophages. Metabolic analysis was explored in further detail below.

**Table 1.**
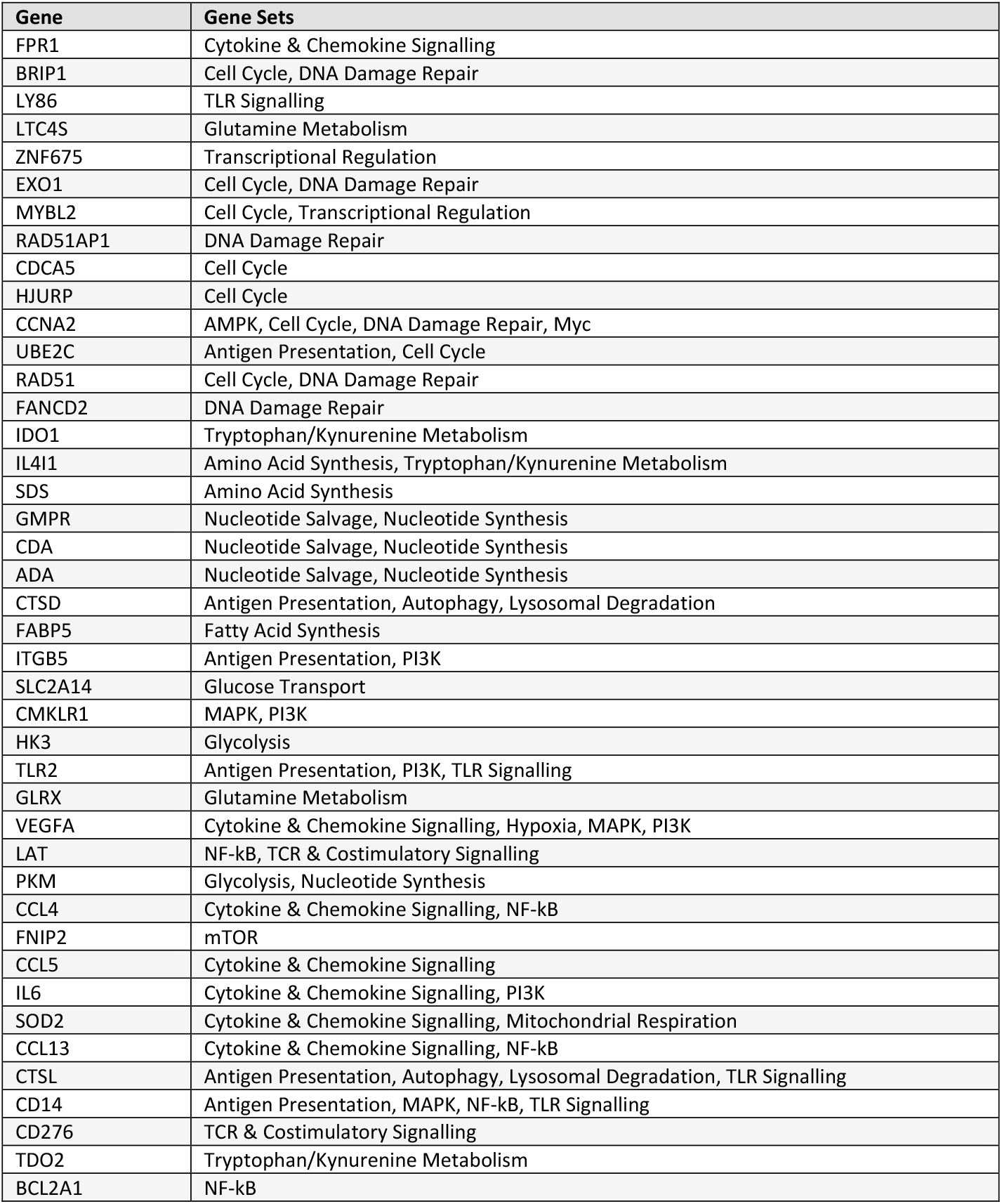
Genes Most Differentially Expressed Between Treatment Groups. Those genes found to be most significantly differentially expressed between the treatment groups are shown with their respective gene sets.

**Figure 1.**
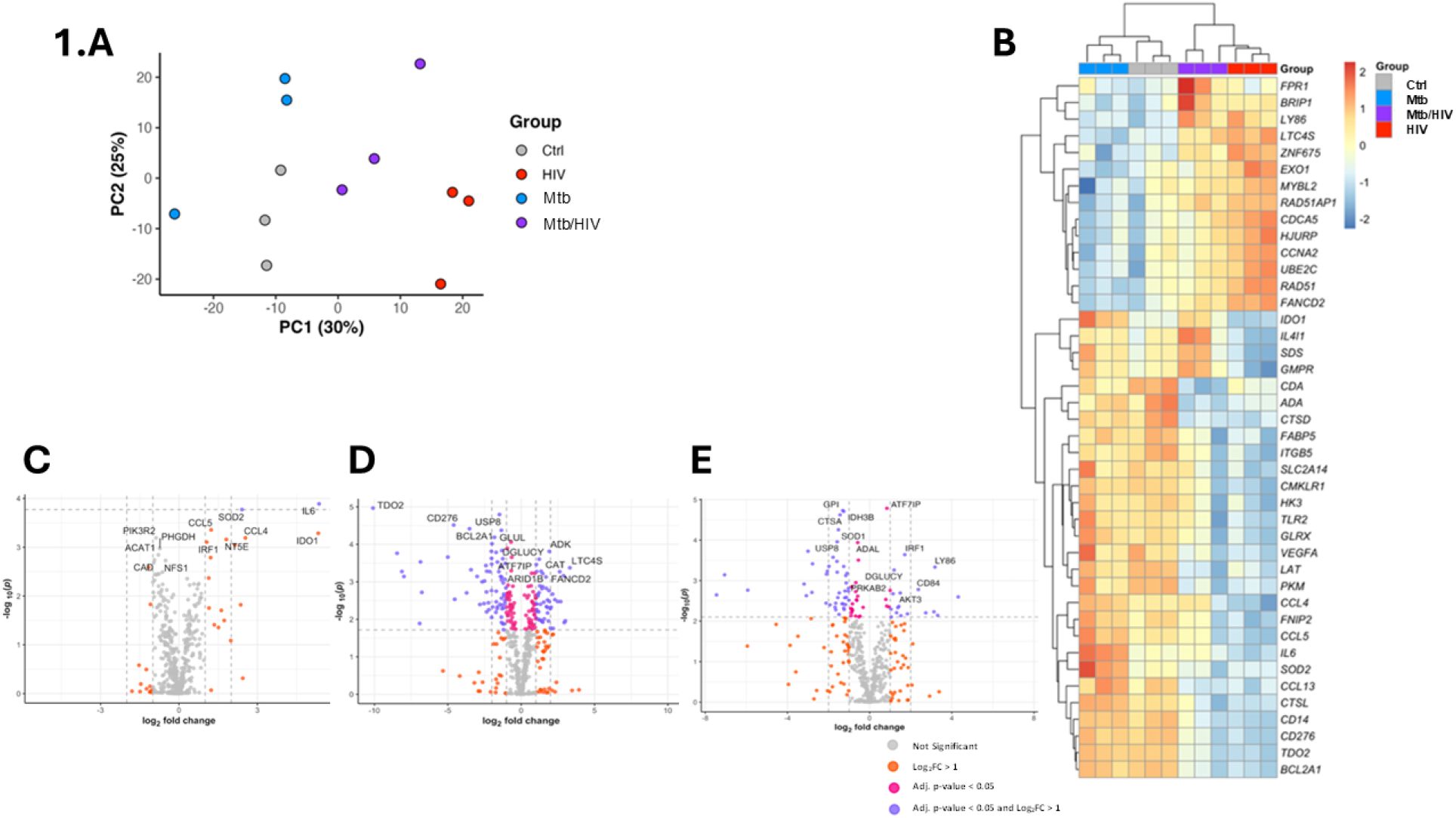
Transcriptional Signatures of Mtb-, HIV-, and Mtb/HIV Co-infected Human Macrophages. Human U937 and U1 macrophages were infected with virulent H37Rv Mtb and mRNA lysates were processed on the NanoString® nCounter® platform Metabolic Pathways panel. Data were analysed with NanoString® nSolver™ 4.0 Advanced Analysis Software, using in-built QC, housekeeping and normalisation processes. A: All treatment groups clustered separately by principal component analysis, suggesting that gene expression changes differentiated the treatment groups. B: Those genes found to be most differentially expressed across the treatment groups are shown in the heatmap, and treatment groups can be seen to cluster separately by hierarchical clustering. C-E: The volcano plots demonstrate gene expression changes in individual genes in Mtb-, HIV-, and Mtb/HIV co-infected macrophage, respectively. Those statistically significant transcriptional changes of greatest magnitude (log2FC > 1) are indicated by purple dots. Pink dots denote statistically significant changes of lower magnitude (log2FC < 1), and orange dots indicate non-significant changes of high magnitude (log2FC > 1). Remaining non-significant changes are coloured in grey.

**Figure 2.**
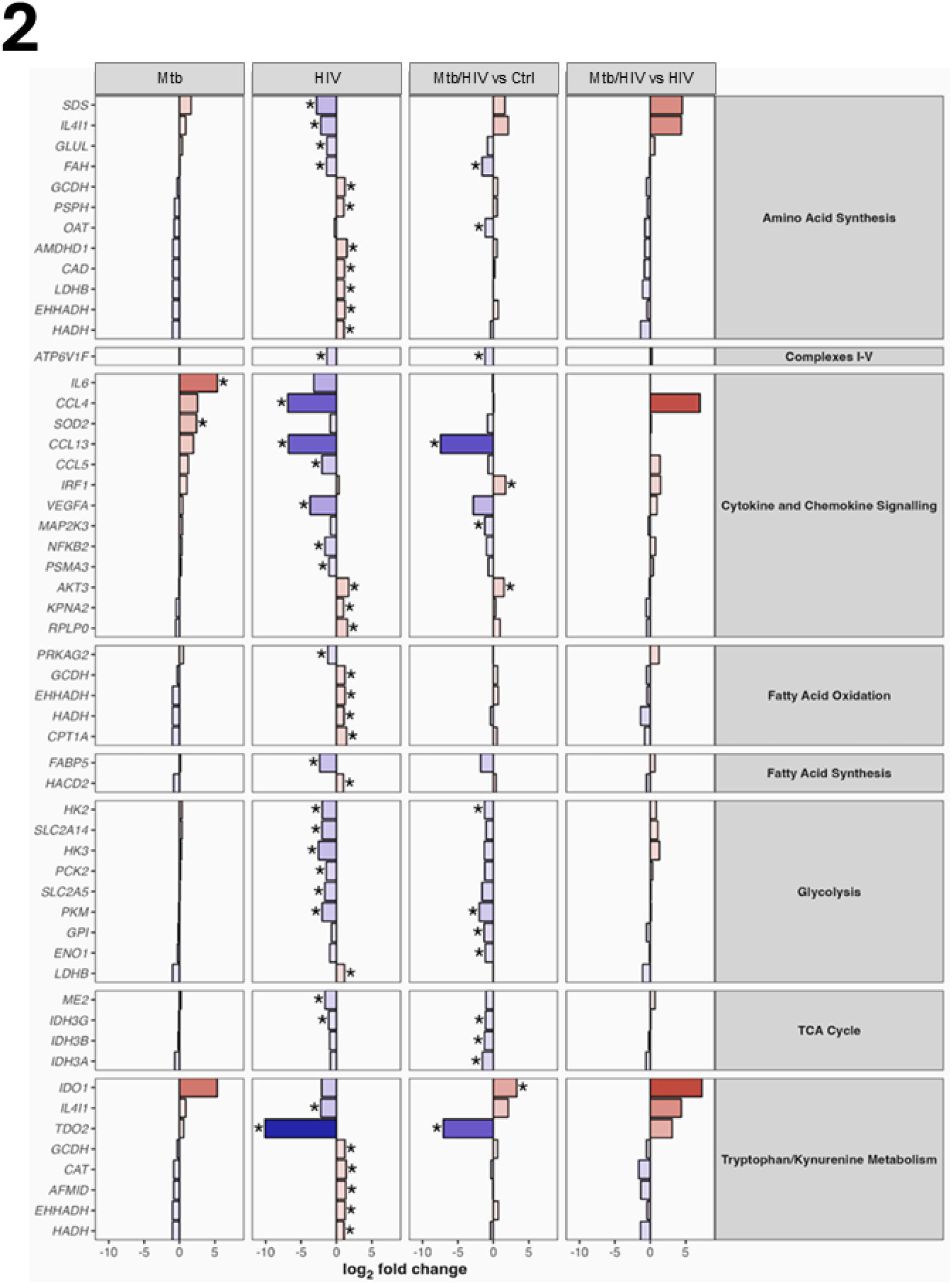
Specific pathways of interest were identified and compared across the treatment groups. Human U937 and U1 macrophages were infected with virulent H37Rv Mtb and mRNA lysates were processed on the NanoString® nCounter® platform Metabolic Pathways panel. Data were analysed with NanoString® nSolver™ 4.0 Advanced Analysis Software, using in-built QC, housekeeping and normalisation processes. Significant gene expression changes of high magnitude (log2FC > 1) within specific pathways of interest were identified and compared across the treatment groups. P-value adjustment for false discovery rate (FDR) was calculated using Benjamini-Hochberg (BH) method, and statistical significance expressed as adj. p-value (q-value), and defined as *q < 0.05.

### Mtb/HIV Co-infected Human Macrophages do not Express a Glycolytic Transcriptional Profile

“Mtb” and “HIV” infected groups induced markedly different glycolytic gene expression profiles (Table S1 and S2, Fig. 3A). Mtb infection did not induce significant changes in glycolytic gene expression, however trends towards induction of *HK2, HK3* and *SLC2A14*, and reduced expression of *FBP1* and *LDHB* suggested a profile consistent with induced aerobic glycolysis. HIV infection significantly reduced gene expression across nearly all glycolytic genes, including all HK isomers, *PFKL, PKM, GPI* and all glucose transporters (*SLC2A1, SLC2A5* and *SLC2A14*) compared to control. Furthermore, HIV significantly reduced the expression of *LDHA* (which primarily catalyses the conversion of pyruvate to lactate), but significantly induced *LDHB* (catalyses lactate to pyruvate), suggesting that HIV-infected human macrophages exhibit a transcriptional profile consistent with reduced glycolysis. Changes in glycolytic gene expression were noted to be of greater significance and magnitude in HIV-infected macrophages compared to Mtb-infected macrophages (Figure 3A).

**Figure 3.**
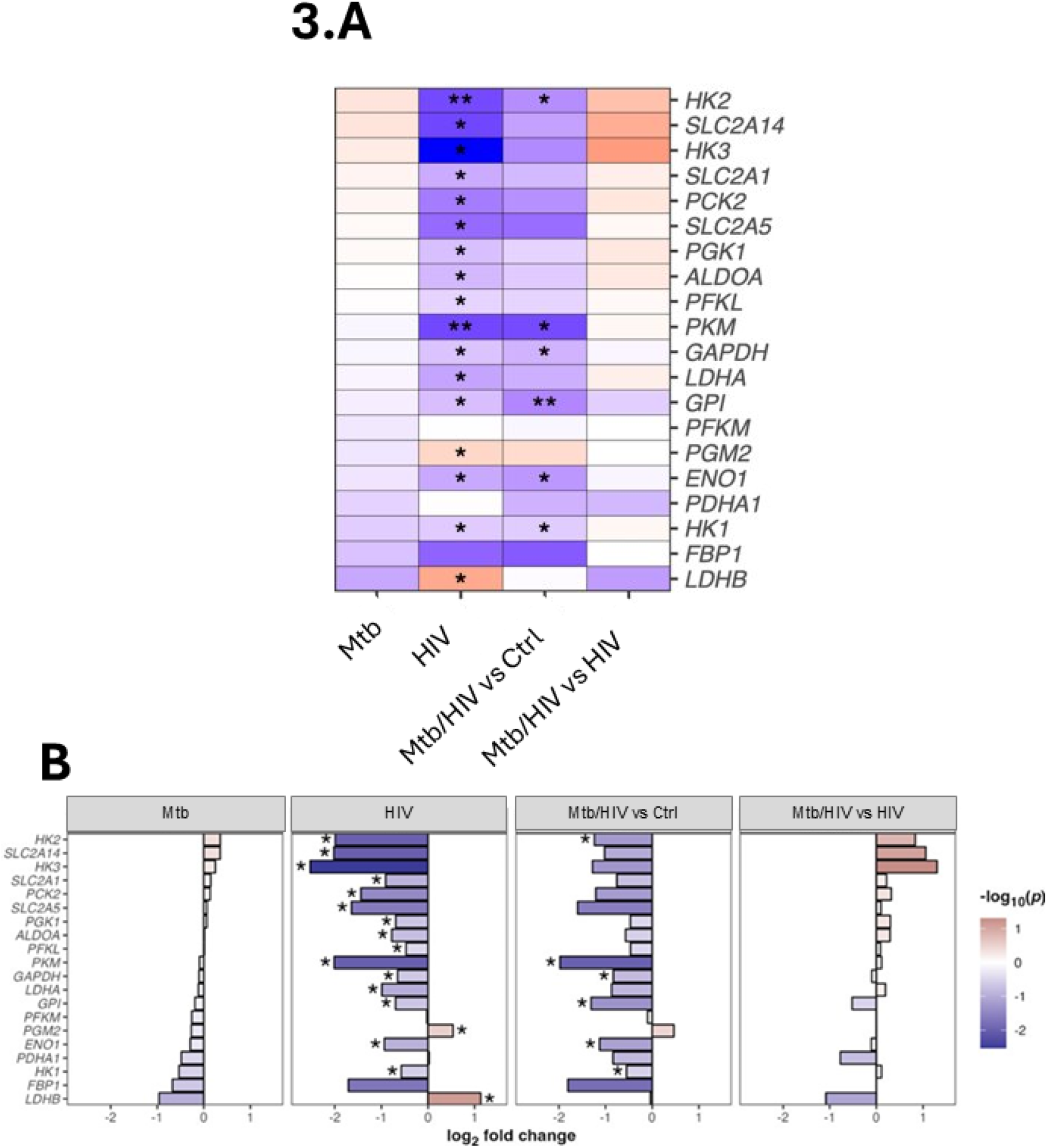
Reduced expression of Glycolytic Genes in Mtb and HIV/Mtb-infected Human Macrophages. Human U937 and U1 macrophages were infected with virulent H37Rv Mtb and mRNA lysates were processed on the NanoString® nCounter® platform Metabolic Pathways panel. Data were analysed with NanoString® nSolver™ 4.0 Advanced Analysis Software. Gene pathways were assigned by NanoString®. A: Gene expression changes across the “Glycolysis” gene set are illustrated for each of the treatment groups, with those significantly differentially expressed indicated by asterisk. Gene expression changes are shown in descending order of log2FC value as per the “TB” treatment group. B: Comparative log2FC bar chart for each of the treatment groups allows visual comparison of global gene expression changes within the gene set. The “HIV” treatment group exhibited a transcriptional profile of markedly reduced glycolytic gene expression, which was sustained in the “Mtb/HIV” treatment group, indicating that HIV inhibited Mtb-induced glycolytic gene expression. Data analysis was performed using the NanoString® nSolver™ 4.0 Software Advanced Analysis module, and data visualisations were generated using R programme. P-value adjustment for FDR was calculated using BH method, and statistical significance defined as *q < 0.05, **q < 0.01.

Mtb/HIV co-infected macrophages exhibited a transcriptional profile consistent with reduced glycolytic gene expression (Figure 3A). When compared with control macrophages (“Mtb/HIV vs Ctrl”), Mtb/HIV co-infected macrophages mirrored the gene expression profile of HIV-infected macrophages, suggesting HIV induced strong and consistent changes in the glycolytic gene set (Figures 3A and 3B). However, when controlling for chronic HIV infection (“Mtb/HIV vs HIV”), Mtb appeared to induce a similar glycolytic profile to that observed in Mtb-infected macrophages (“Mtb”) (Figures 3B). These results suggest that Mtb infection induced glycolysis in HIV-infected macrophages, but as HIV-induced changes in glycolytic gene expression were of greater magnitude, these changes were not observed in Mtb/HIV co-infected macrophages when compared to uninfected control.

### HIV Drives Increased Expression of the Mitochondrial OXPHOS Gene Network and Influences its Transcriptional Profile in Mtb/HIV Co-Infected Macrophages

Mtb infection appeared to restrict mitochondrial OXPHOS gene expression in human macrophages compared to the control, although changes were not found to be statistically significant (Table S3, Figures 4A). Reduced expression was observed across all five ETC complexes, and the “Mtb” treatment group clustered separately from other conditions, as demonstrated in the heatmap in Figure 4B. A profile of unanimously reduced expression was evident from the treatment groups comparative bar chart in Figure 4.C. HIV induced a markedly different mitochondrial OXPHOS transcriptional profile vs. control. Of greatest interest was the almost universally increased expression of Complex I subunits, with similar expression patterns seen for Complexes IV and V (Figure 4.A). Reduced expression of Complex II was consistent with reduced SDH expression.

**Figure 4.**
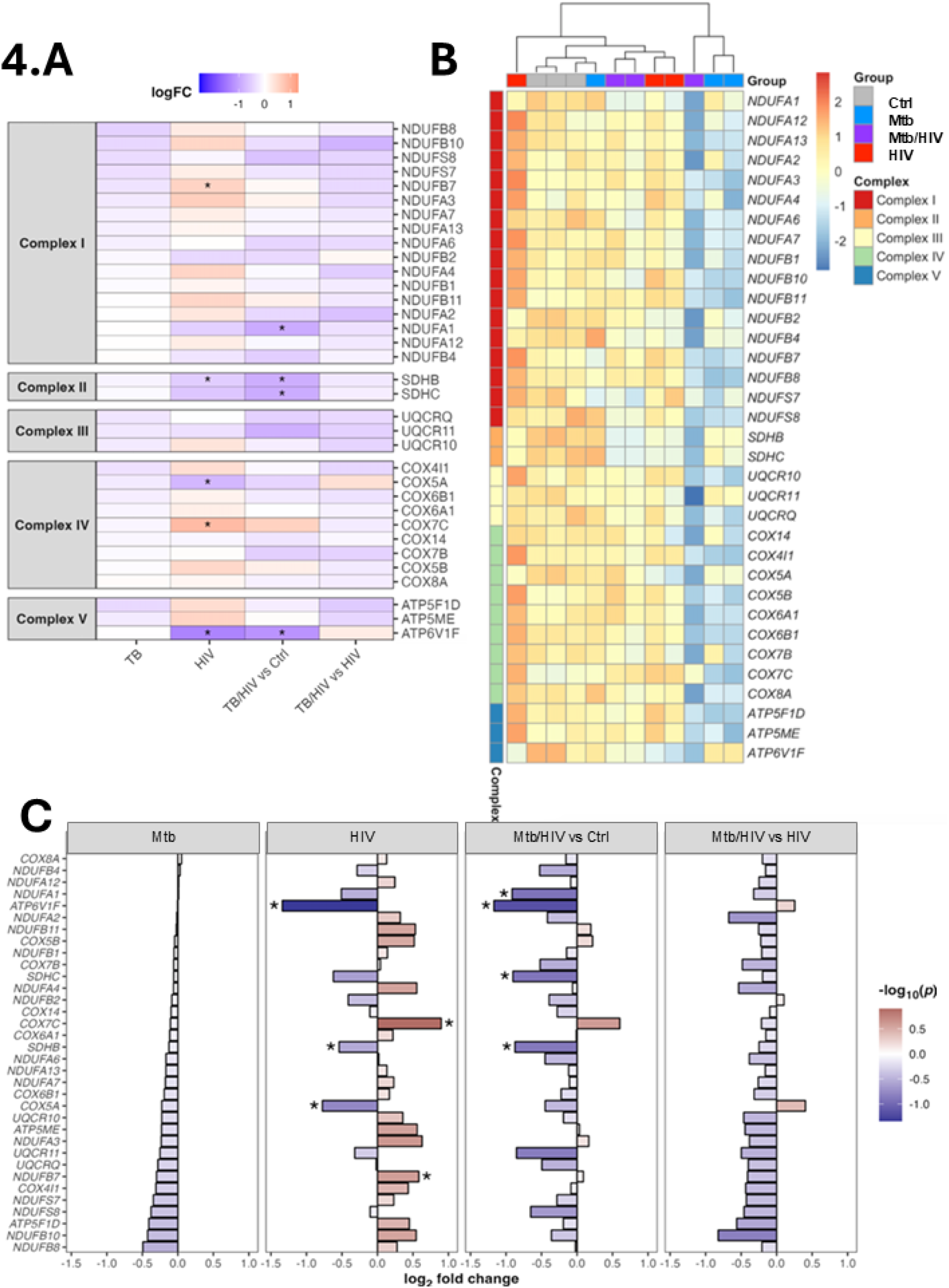
HIV Drives Mitochondrial OXPHOS Gene Expression. Human U937 and U1 macrophages were infected with virulent H37Rv Mtb and mRNA lysates were processed on the NanoString® nCounter® platform Metabolic Pathways panel. Data were analysed with NanoString® nSolver™ 4.0 Advanced Analysis Software. Gene pathways were assigned by NanoString®. A: Gene expression changes across mitochondrial electron transport chain genes from the “Mitochondrial Respiration” gene set are illustrated for each of the treatment groups, with those significantly differentially expressed indicated by asterisk. Gene expression changes are shown in ascending order of log2FC value as per the “TB” treatment group. B: Gene expression heatmap demonstrated universally restricted mitochondrial OXPHOS gene expression in the “Mtb” treatment group, whilst “HIV” and “Mtb/HIV” groups clustered together. C: Comparative log2FC bar chart for each of the treatment groups allows visual comparison of global gene expression changes within the gene set. HIV drove mitochondrial OXPHOS gene induction, with a transcriptional profile markedly different from the “TB” treatment group. Some HIV-induced changes were sustained in the “TB/HIV” treatment group, suggesting that HIV infection disrupted Mtb-induced reduction in mitochondrial OXPHOS gene expression. Data analysis was performed using the NanoString® nSolver™ 4.0 Software Advanced Analysis module, and data visualisations were generated using R programme. P-value adjustment for FDR was calculated using BH method, and statistical significance defined as *q < 0.05.

Interestingly, despite this marked HIV-driven induction in mitochondrial OXPHOS gene expression, “Mtb/HIV” macrophages still exhibited overall reduced mitochondrial OXPHOS transcriptional profiles (Figure 4A). The “Mtb/HIV” treatment group, however, clustered separately from the “Mtb” treatment group, suggesting that HIV co-infection still influenced the transcriptional profile of Mtb/HIV co-infected macrophages. As can be seen in Figure 4C, HIV-infected macrophages exhibited a markedly different transcriptional profile compared to the other treatment groups.

### HIV gp120 stimulation disrupted early Warburg metabolism in Mtb-infected human MDM through the induction of oxidative metabolism

HIV gp120-treated MDM were analysed on the Seahorse XFe24 Analyser using the Acute Injection protocol described in the methods section. After four resting OCR/ECAR baseline measurements, all wells received an injection of either iH37Rv (Experimental) or Seahorse XF RPMI medium (Control) at 28 minutes and OCR/ECAR were measured for a total of 180 minutes, as shown in Figures 5A – 5C. Time of iH37Rv injection is indicated and labelled on the graph. Time of OCR and ECAR analysis was chosen as the time of maximum OCR and ECAR response in “Mtb” MDM, and is also indicated on the graph. “TB” MDM (blue) exhibited immediate Warburg metabolism following Mtb injection, characterised by significantly increased ECAR (Figures 5A and 5D) and significantly decreased OCR (Figures 5A and 5E), compared to “Control” MDM (grey). “Lo HIV/Mtb” MDM (purple) exhibited a significantly lower ECAR response after Mtb injection, compared to “Mtb” MDM (Figures 5B and 5D). Neither “Lo HIV/Mtb” nor “Hi HIV/Mtb” (red) exhibited any significant change in OCR compared to “Control”, and both exhibited significantly increased OCR compared to “TB” (Figures 5B, 5C, 5E). Phenotypically, both “Lo HIV/Mtb” and “Hi HIV/Mtb” exhibited more energetic profiles, characterised by an increase in both ECAR and OCR (Figures 5G,5H), which contrasted with the Glycolytic profile expressed by “Mtb” MDM, characterised by an increased ECAR and decreased OCR (Figure 5D,E). These results reinforce the notion that Mtb-infected MDM exhibit early Warburg metabolism(33), and demonstrate how this Warburg metabolism is prevented by prior HIV gp120 treatment. Given that of the changes observed, markedly increased OCR was the predominant finding in both “Lo HIV/Mtb” and “Hi HIV/TB” treatment groups compared to “Mtb” MDM, this suggested that HIV gp120 stimulation of MDM rewired mitochondrial metabolism. Mitochondrial function was therefore examined using the Mito Stress Test to assess whether HIV gp120 treatment impaired specific mitochondrial kinetics during Mtb infection.

**Figure 5.**
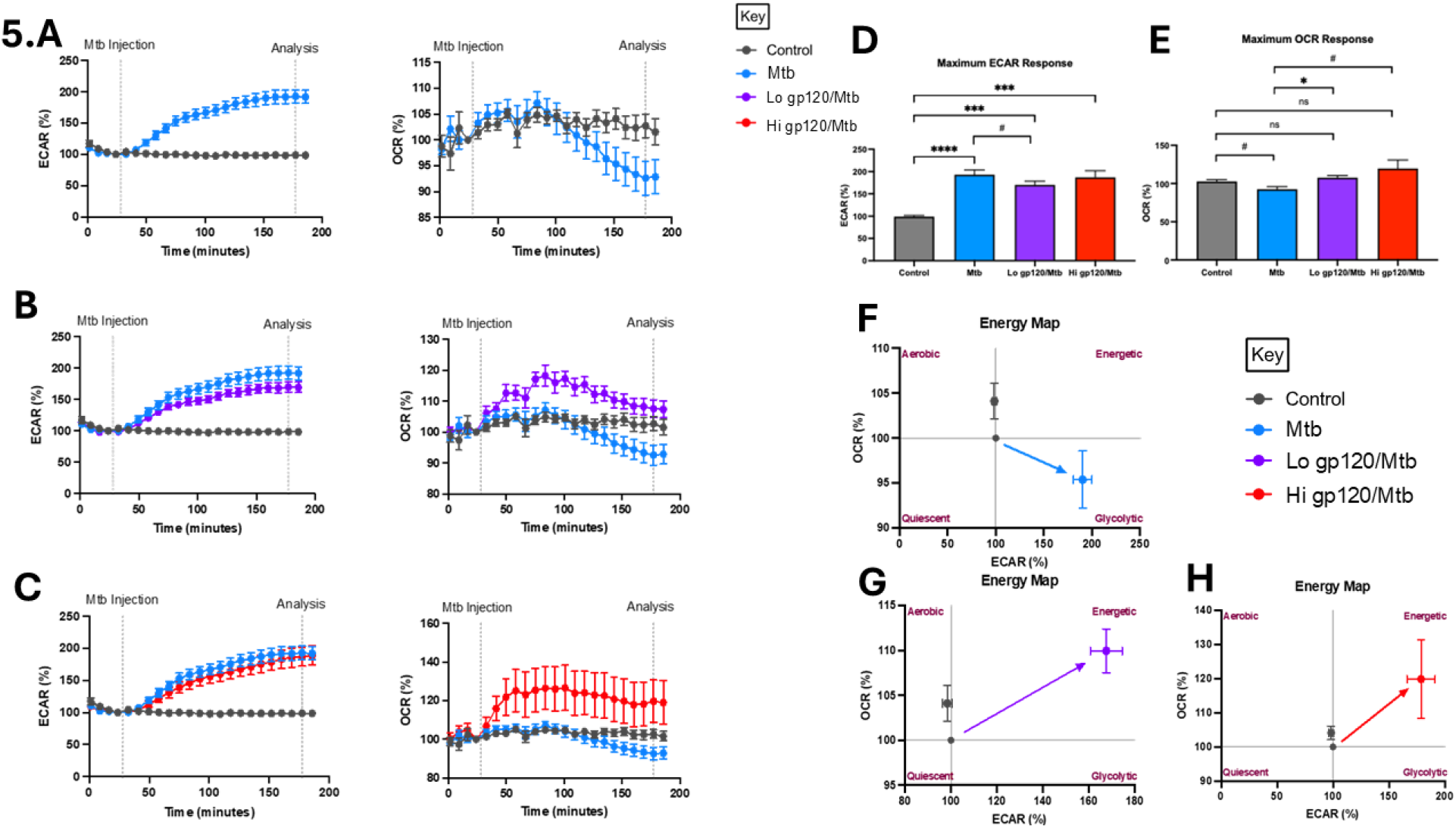
HIV gp120 treatment prevents early Warburg metabolism in human MDM. MDM were treated with either low (100 ng/mL) or high (5,000 ng/mL) dose HIV gp120 or media control for 24 hr before analysing on the Seahorse XFe 24 Analyser. After four basal measurements, samples received an injection of iH37Rv (blue, red, purple) or media control (grey) at a run time of 28 minutes, as indicated on the graph (“Mtb Injection”), and changes in ECAR/OCR were then measured in real-time for a total of 180 minutes. MDM supernatants were collected at the end of the run. For statistical analyses a single time point was used for comparisons. This time point was selected based on the maximum ECAR and OCR changes exhibited by the “Mtb” MDM (blue) group, and is also indicated on the graph (“Analysis”). “Mtb” MDM immediately increased ECAR and decreased OCR after receiving iH37Rv injection, consistent with Warburg metabolism (A). “Lo gp120/Mtb” (purple) and “Hi gp120/Mtb” (red) MDM also immediately increased both ECAR after iH37Rv injection and OCR (B and C). All treatment groups exhibited a significantly increased ECAR compared to “Control” MDM (grey), however the magnitude of ECAR induction in “Lo gp120/Mtb” MDM was significantly lower than in “Mtb” MDM (D). “Mtb”MDM significantly decreased OCR compared to “Control” MDM, and neither “Lo gp120/Mtb” nor “Hi gp120/Mtb” MDM demonstrated any significant change in OCR compared to “Control” MDM. Both “Lo gp120/Mtb” and “Hi gp120/Mtb” MDM exhibited significantly increased OCR compared to “Mtb” MDM (E). “Mtb” MDM expressed a glycolytic phenotype (F), whereas both “Lo gp120/Mtb” and “Hi gp120/Mtb” MDM expressed energetic phenotypes (G and H). Data are presented as mean and standard error of the mean (SEM) (n = 8 biological replicates). Analysis by one-way ANOVA (*p < 0.05, **p < 0.01, ***p < 0.001, ****p < 0.0001) or paired Student’s t-test (#, p < 0.05).

### HIV gp120 Treatment Reprogrammed Mitochondrial Function in Mtb-infected Human MDM

Our transcriptional data indicated that chronically HIV-infected human macrophages exhibit increased expression of mitochondrial ETC genes, and the findings above suggest HIV gp120 treatment of MDM induces a metabolic phenotype characterised by high oxygen consumption during Mtb infection instead of Warburg metabolism. It is therefore plausible to suggest that Mtb-infected MDM mitochondrial function might be disturbed by HIV gp120 treatment. This hypothesis is supported by previous reports of HIV gp120-induced disturbances of mitochondrial function (32, 34).

Neither the “Lo gp120” nor “Hi gp120” treatment group significantly impaired any parameters of mitochondrial function (Fig.S1). Mitochondrial function was then investigated in Mtb/HIV gp120 co-infection. MDM were treated with HIV gp120 as before, and were then infected with iH37Rv Mtb 30 – 60 minutes before Mito Stress Test. “Mtb” MDM showed no changes in OCR (Figure 6A and B), however significantly increased total ECAR (Figures 6C and 6D). Both “Lo gp120/Mtb” and “Hi gp120/Mtb” MDM exhibited impaired capacity to increase ECAR, compared to “Mtb” MDM (Figures 6D). This was most significant in the “Hi gp120/Mtb” treatment group, where no statistical difference in total ECAR was found compared to “Control” MDM (Figure 6D). “Lo gp120/Mtb” MDM had significantly reduced SRC compared to both “Control” and “Mtb” MDM. A similar trend was observed in “Hi gp120/Mtb” MDM, although this change was not found to be statistically significant (p = 0.08) (Figure 6E). This finding indicates an exacerbated effect on SRC during co-stimulation over that observed in HIV gp120-treated MDM without Mtb infection.

**Figure 6.**
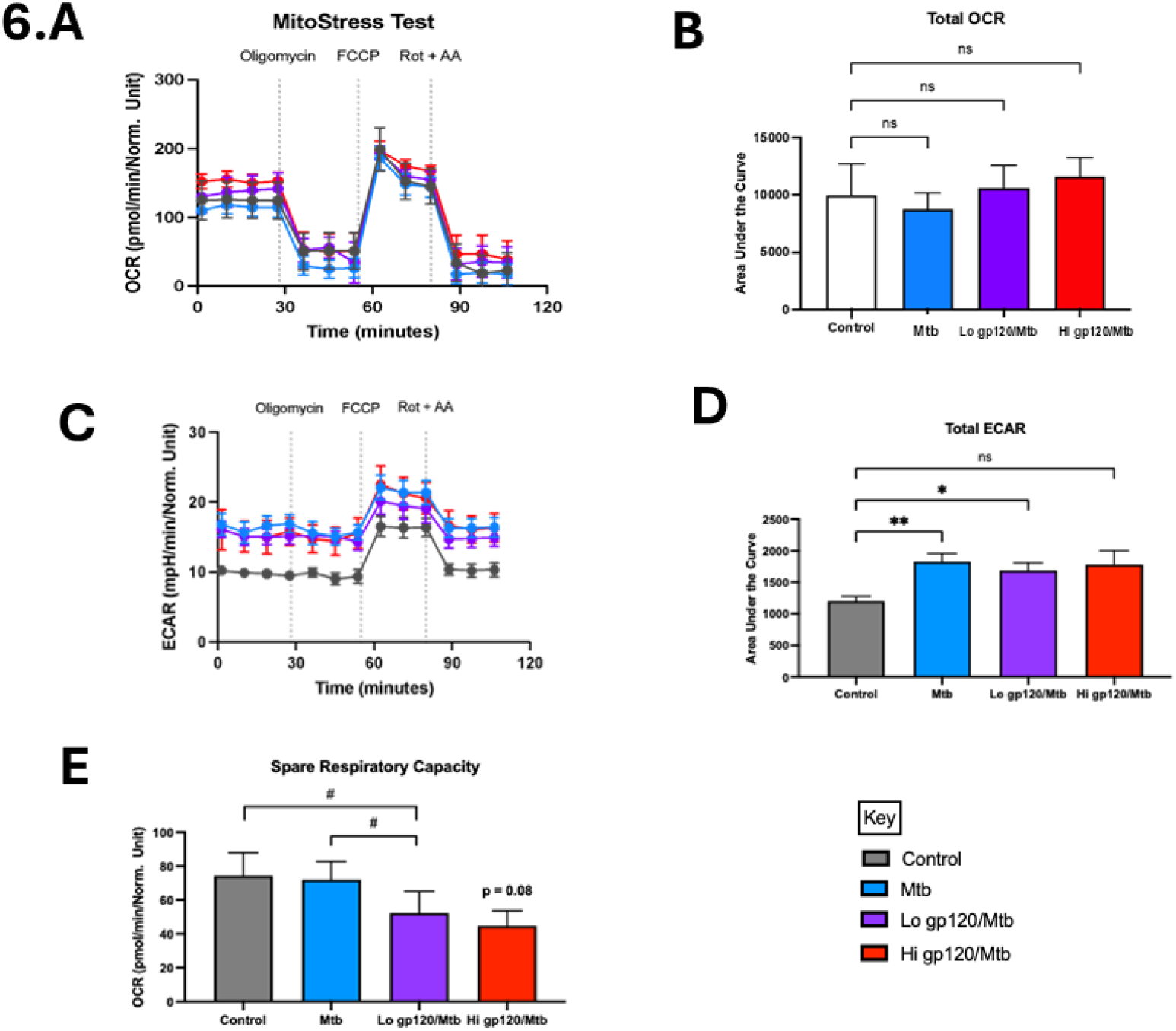
HIV gp120 treatment rewired mitochondrial function in Mtb-infected MDM. MDM were treated with either low (100 ng/mL) or high (5,000 ng/mL) dose HIV gp120 or media control for 24 hr. MDM were infected with iH37Rv infection at an MOI of ~70% infectivity and 5-10 bacilli/cell before analysing on the Seahorse XFe 24 Analyser, using the MitoStress Protocol. Total OCR showed no change across treatment groups (A,B). Total ECAR was significantly increased in Mtb and “Lo HIV/TB” only compared to compared to “Control” MDM(C, D). “Lo gp120/Mtb” MDM had significantly reduced Spare Respiratory Capacity compared to “Mtb” and “Control” MDM, although no statistically significant difference in Spare Respiratory Capacity was observed in “Hi gp120/Mtb” MDM compared to “Mtb” or “Control” MDM (E). p-values and statistical tests used. p-values, n number, mean and SEM, and statistical tests used. Data are presented as mean and SEM (n = 5 biological replicates). Analysis by one-way ANOVA (*p < 0.05, **p < 0.01, ***p < 0.001, ****p < 0.0001) or paired Student’s t-test (#, p < 0.05).

### HIV gp120 reduced TNF production in human MDM infected with Mtb

Considering Mtb-induced glycolytic reprogramming is an established paradigm in human macrophage host defence, that has been specifically linked to downstream effector functions including IL-1β secretion and bacillary killing (35), interruption of this early Warburg metabolism could be considered a mechanistic functional impairment of macrophage host defence. Consequently, effector functions were investigated as part of this Mtb/HIV co-infection model assessment to strengthen the findings of impaired Mtb-induced macrophage metabolism described above.

Firstly, cytokines of known importance in Mtb infection, and/or those recognised to be under metabolic influence, were measured and included TNFα, IL-1β and IL-10. Macrophage supernatants were collected following bioenergetic analysis, and secreted cytokines were measured by sandwich ELISA. As expected, “Mtb” MDM secreted significantly more TNFα compared to “Control” MDM. No difference was observed between “Mtb” MDM and “Lo HIV/Mtb” MDM, however “Hi gp120/Mtb” MDM failed to significantly induce TNFα secretion (Figure 7A). “Mtb” MDM did not induce significantly more IL-1β compared to “Control” MDM, and no differences in IL-1β secretion were observed between any of the treatment groups (Figure 7A). Similarly, “Mtb” MDM did not significantly induce IL-10 secretion, nor were there any changes detected between any of the treatment groups (Figure 7A). Macrophage TNFα secretion has been shown to peak 2 – 4 hr following LPS stimulation (36), however it is unlikely that macrophage IL-1β or IL-10 secretion would have increased significantly at this time point.

**Figure 7.**
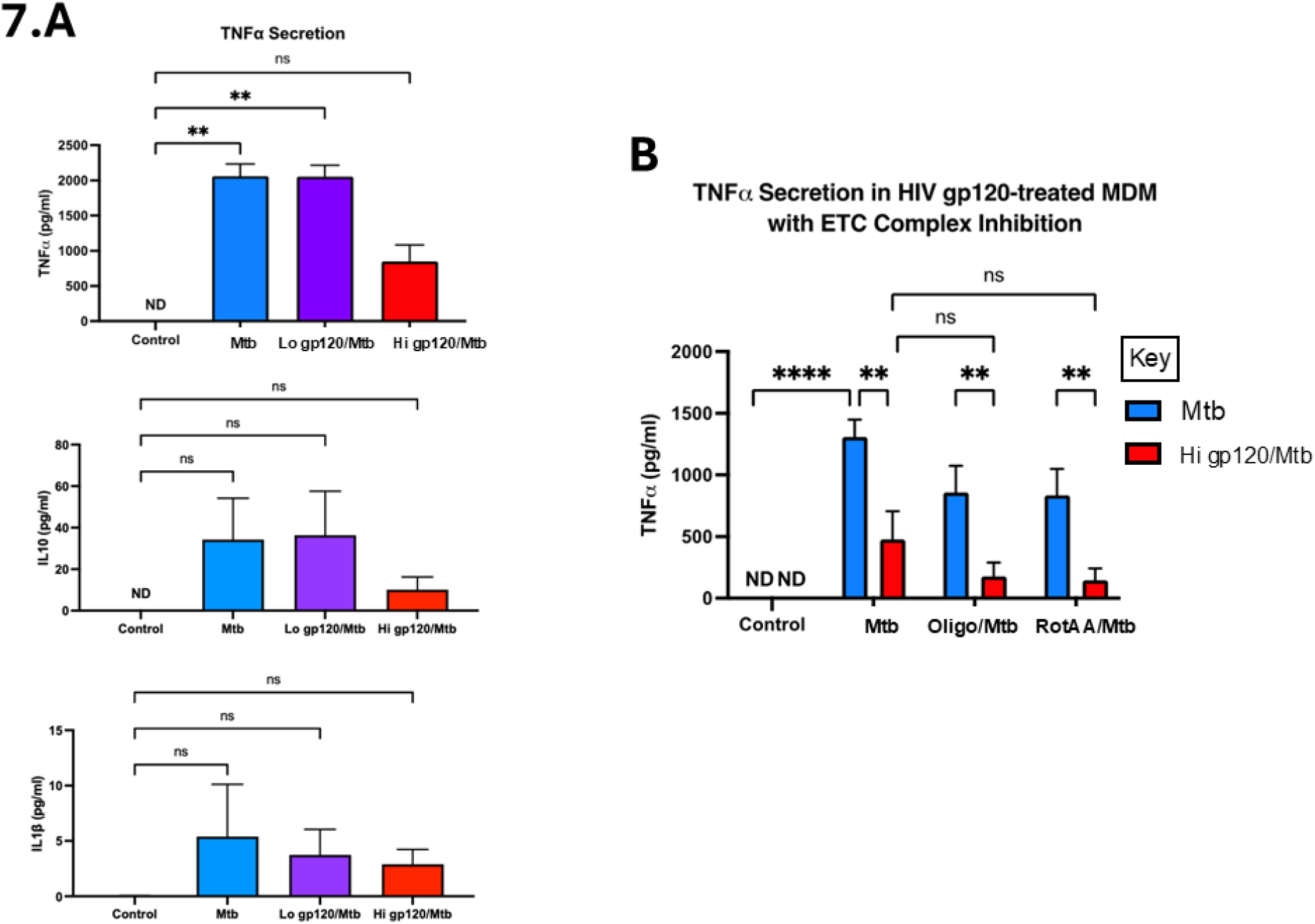
High dose HIV gp120 treatment reduced TNFα secretion in Mtb-infected human MDM which is dependant the ETC. MDM were treated with low or high dose HIV gp120 or media control for 24 hr. MDM were then infected with iH37Rv infection at an MOI of ~70% infectivity and 5-10 bacilli/cell. Supernatants were collected at 3 hr post infection and analysed by sandwich ELISA for cytokine concentrations. “Mtb” MDM secreted significantly higher concentrations of TNFα compared to “Control” MDM. “Hi gp120/Mtb” MDM secreted significantly lower concentrations of TNFα compared to “Mtb” MDM. No difference in IL-1β and IL-10 secretion was detected between the treatment groups (A). MDM were treated with high dose HIV gp120 or media control for 24 hr before analysing on the Seahorse XFe 24 Analyser, using the Acute Injection protocol. Samples received an injection of either medium control (“Control”, “Mtb”), oligomycin (“Oligo/Mtb”) or rotenone/antimycin A (“RotAA/Mtb”), followed by an injection of either medium control (“Control”) or iH37Rv Mtb (“Mtb”, “Oligo/Mtb”, “RotAA/Mtb”). Supernatants were collected at 3 hr post Mtb injection and TNFα measured using sandwich ELISA. “Mtb” MDM secreted significantly more TNFα compared to “Control” MDM. “Hi gp120/Mtb” MDM secreted significantly lower concentrations of TNFα compared to “Mtb” MDM, and TNFα concentrations were not restored by prior treatment with oligomycin (“Oligo/Mtb”) or rotenone/antimycin A (“RotAA/Mtb”) (B). Data are presented as mean and SEM (n = 4 biological replicates). Analysis by two-way ANOVA (*p < 0.05, **p < 0.01, ***p < 0.001, ****p < 0.0001).

### Reduced TNFα secretion from Mtb-infected human MDM is linked to impaired mitochondrial function

The data presented so far indicates that HIV gp120 treatment prevents Warburg metabolism in Mtb-infected MDM and, at high dose, restricts TNFα secretion. Whether these findings are related, and whether macrophage metabolism might exert control over TNFα secretion, is unclear. Considering the most marked difference between gp120/Mtb MDM and Mtb MDM was the rate of oxygen consumption, and that HIV gp120 treatment significantly impaired macrophage SRC, it was hypothesised that TNFα secretion might depend on mitochondrial function. To determine whether the observed increase in OCR was responsible for reduced TNFα secretion in HIV gp120-treated Mtb-infected MDM, macrophage mitochondrial OXPHOS was inhibited, using specific ETC complex inhibitors, prior to Mtb infection. TNFα concentration was then again measured in MDM supernatants post bioenergetic flux analysis to assess whether inhibition of mitochondrial OXPHOS might restore TNFα secretion. As before, MDM were either treated for 24 hr with HIV gp120 or left untreated. Again, only high dose HIV gp120 was included, as this dose inhibited TNFα secretion. After 24hr of HIV gp120 treatment, MDM were processed on the Seahorse Analyser and received two injections. After 28 minutes, MDM received a first injection of either oligomycin (ATP synthase inhibitor), R/AA (Complex I/Complex III inhibitors) or Seahorse XF RPMI medium (control). After a further 28 minutes, MDM then received a second injection of either iH37Rv Mtb or Seahorse XF RPMI medium (control). ECAR and OCR were then measured for a total of 180 minutes and supernatants were collected at the end of the analysis, which were then processed by sandwich ELISA.

As before, hi gp120/Mtb MDM had significantly reduced TNFα secretion compared to Mtb MDM (Figure 7B). Both oligomycin and R/AA treatment appeared to decrease Mtb MDM TNFα secretion, as indicated by the blue bars, although this reduction was not found to be statistically significant. Additionally, neither oligomycin nor R/AA restored TNFα secretion in hi gp120/Mtb MDM, but in fact appeared to further reduce TNFα secretion, although this was not found to be statistically significant.

## Discussion

Macrophage metabolism during HIV infection is poorly defined, however, it is well established that HIV targets cells with heightened metabolism for infection, that helps enable efficient viral replication and virion assembly (2, 3, 5, 6). HIV-induced changes in cell metabolism has been characterised in CD4^+^ T cells (10, 37–41), with similar findings reported in monocytes (42, 43). However, observations of HIV-induced changes in macrophage metabolism are limited to a small number of studies (5, 7, 8, 10, 11), and altered glutamine metabolism remains the only reproducible finding (7, 8, 11).

Macrophage immunometabolism in Mtb/HIV co-infection has not been previously defined, despite the central role of metabolism in orchestrating an effective anti-tuberculous immune response and the new opportunities it might represent for therapeutic targeting. This study aimed to characterise the metabolic transcriptional profile of HIV-infected human macrophages, and to define specific metabolic pathway perturbations that might be important in corrupting macrophage immunity in Mtb infection thereby promoting TB disease.

Mtb-infected macrophages expressed a transcriptional profile characterised by induced HK2/3 and glucose transporters, with reduced expression of LDHB and the gluconeogenic FBP1. Conversely, HIV infection clearly suppressed the macrophage glycolytic profile, characterised by significantly reduced expression of *HK2, HK3, GLUT5, GLUT14, PKM* and *PCK2*, with various other genes trending toward reduced expression, and significantly increased expression of LDHB. Mtb/HIV co-infection induced a transcriptional profile similar to that observed in HIV-infected macrophages, suggesting that HIV infection inhibited Mtb-induced macrophage glycolysis. These findings are in keeping with observations of reduced glycolysis in HIV-infected macrophages(7, 10), and represent the first investigation of the glycolytic transcriptional profile in Mtb/HIV co-infected macrophages. Mtb-infected macrophages appeared to inhibit mitochondrial OXPHOS gene expression across all five ETC complexes. By contrast, HIV infection appeared to drive mitochondrial OXPHOS gene expression, with significantly induced expression observed in complexes I and IV. Mtb/HIV co-infected macrophages generally maintained a transcriptional profile of reduced mitochondrial OXPHOS, similar to Mtb-infected macrophages. However, HIV-induced increased expression of subunits of mitochondrial complexes I and IV were sustained in Mtb/HIV co-infected macrophages. Inhibition of Complex I has been explored therapeutically through the use of Metformin as an adjuvant therapy in TB disease (44, 45). Considering this, and its central role in NAD^+^ recycling, this finding could be of particular relevance in Mtb/HIV co-infection.

The HIV envelope protein, gp120, has been shown to influence cellular metabolism in various human models. HIV gp120-treated *S. pneumoniae*-infected MDM exhibit reduced proton leak with maintained mitochondrial membrane potential, and impaired pneumococcal killing (32). HIV gp120 treatment of human glioma cells induces glycolysis (46) and HIV gp120-treated neurons exhibit reduced SRC compared to untreated control neurons in rat brains (34). Persistent circulating HIV gp120 has been observed in the serum of PLWH who are taking virally suppressive ART (47), and has also been detected in human BAL fluid (32) and lung tissue (48). Whether this circulating HIV gp120 might exert an inhibitory influence over the capacity of tissue-resident AM, or indeed migratory MDM, to effectively respond to Mtb during the critical early phases of infection has not been previously investigated, and might have potentially significant consequences for PLWH on effective ART.

The results presented here explore the early metabolic changes in human MDM in a Mtb/HIV co-infection model, and demonstrate how HIV gp120 stimulation of MDM corrupts immediate Mtb-induced glycolytic reprogramming, or Warburg metabolism. HIV gp120-treated MDM do not shift metabolism from mitochondrial respiration towards a glycolytic phenotype, as measured by bioenergetic analysis, when challenged with Mtb stimulation. Instead, HIV gp120-treated Mtb-infected MDM express an “energetic” phenotype, characterised by high rates of both oxygen consumption and extracellular acidification. Furthermore, HIV gp120-treated Mtb-infected MDM have significantly reduced SRC compared to Mtb-infected MDM, indicating an impaired capacity to respond to infection. These findings align with previous observations of HIV gp120-induced reduction in SRC (34), and impaired mitochondrial kinetics in an HIV gp120/Pneumococcus co-infection model (32). It has also been observed that Mtb-infected smoker’s AM have reduced SRC compared to non-smokers’ AM (49), and Mtb-infected cord blood macrophages trend towards reduced SRC compared to adult MDM (50), suggesting that SRC is an important indicator of impaired macrophage metabolism in various other TB “at risk” groups. Together these results indicate that HIV gp120 treatment of MDM prevents early Mtb-induced Warburg metabolism and may impair their capacity to respond to insult. Such effects could reasonably be suggested to lead to long term immunometabolic paralysis of HIV gp120-exposed immune cells, corrupting their early anti-Mtb response. Interestingly, this result was true for both “Lo gp120/Mtb” and “Hi gp120/Mtb” treatment groups, suggesting the effect was not dose-dependent and that low dose HIV gp120 was sufficient to prevent glycolytic reprogramming in Mtb-infected MDM. Since the “Lo gp120/TB” treatment group models the HIV gp120 concentrations that have been documented in ART-suppressed HIV (32, 47), this finding is of particular interest when considering the continued risk of TB disease in PLWH who are on virally-suppressive therapy (24, 29).

Mtb-induced TNFα secretion from human macrophages is necessary for bacillary control (51, 52). Mtb/HIV co-infected human macrophages secrete lower concentrations of TNFα compared to Mtb-infected macrophages and this impairs Mtb-induced apoptosis (15). The data presented here demonstrates how high dose HIV gp120 stimulation of MDM suppressed secretion of Mtb-induced TNFα, however a direct link between reduced TNFα secretion and impaired Warburg metabolism was not immediately apparent. Although not traditionally considered to be metabolically regulated, there have been a number of reports linking TNFα secretion to macrophage metabolism(53). Our lab has shown previously that corrupted early macrophage metabolism in Mtb-infected neonatal cord blood macrophages resulted in a significant reduction in TNFα secretion when compared to adult MDM (50). Whilst the data presented in our study did not find a direct link between HIV gp120-induced MDM metabolism and reduced TNFα secretion, inhibition of MDM mitochondrial OXPHOS was found to noticeably and reproducibly restrict Mtb-induced TNFα secretion. Supporting these findings are previous observations of reduced TNFα-associated inflammation in LPS-treated mice after treatment with metformin (a mitochondrial complex I inhibitor) (54, 55), and reduced TNFα secretion from Mtb lysate-stimulated human PBMC after treatment with metformin (56). Intracellular NAD^+^:NADH ratios might therefore govern TNFα secretion capacity, and pharmacological bolstering of cellular NAD^+^ concentrations by the administration of nicotinamide might therefore represent an avenue for host-directed therapy exploration as a cheap and safe adjuvant for the treatment of Mtb/HIV co-infection.

This study employed the use of an Mtb/HIV co-stimulation model in human MDM, rather than virulent Mtb/live HIV co-infected human MDM, which limits the impact of these findings. While using virulent Mtb and moreover, clinical strains is important and warrants further study, these experiments were designed to allow for the assessment of how HIV infection would make an individual more vulnerable to TB, we therefore used a model of attenuated Mtb to model effective early clearance and appropriate immune responses not confounded by virulent Mtb’s ability to modulate metabolism. Nonetheless, this data represents an important first step towards the understanding of immunometabolism in TB/HIV co-infection that can be further built upon with increasingly translational models. Furthermore, examining the effect of HIV gp120 alone is a model that might be relevant to PLWH who are taking suppressive ART (32, 47, 48, 57).

In conclusion, PLWH on virally suppressive therapy continue to experience a significantly greater risk of TB disease than do those without HIV (24–26, 28). Despite an appreciation for immunometabolism in Mtb infection (35, 50, 58, 59), and of the importance of early events in TB disease outcomes (60), we have a limited understanding of the influence of HIV co-infection and its influence on TB disease trajectory (13). These results demonstrate how HIV gp120 treatment prevents early Warburg metabolism in Mtb-infected MDM, and represents the first examination of phenotypic immunometabolism in Mtb/HIV co-infection. How this immunological pathway might be related to HIV-impaired anti-Mtb immunity in human macrophages, and whether it can be therapeutically targeted, will be an important avenue to explore for the future development of novel host-directed immunotherapies.

## Acknowledgments

This work was funded by the Royal Hospital of Dublin City Trust and the Health Research Board (EIA-2019-010, ILP-POR-2019-106). The funders had no role in study design, data collection and interpretation, or the decision to submit the work for publication. Dr. John Browne, Professor Steven Gordon, and the University College Dublin NanoString laboratory staff. We acknowledge the key contributions of The Irish Blood Transfusion Services. The following reagent was obtained through BEI Resources, NIAID, NIH: *Mycobacterium tuberculosis*, Strain H37Rv, Gamma-Irradiated Whole Cells, NR-49098Bei Resources, NIAID, NIH.

## Supplementary Figure Legends

**Figure S1. HIV gp120 treatment alone does not impair mitochondrial function**. MDM were treated with low or high dose HIV gp120 or media control for 24 hr before analysing on the Seahorse XFe 24 Analyser, using the MitoStress Protocol. No differences in total OCR or ECAR were found between any of the treatment groups (A and B), nor were any differences in baseline OCR (C) or baseline ECAR (D).

